# Data-adaptive pipeline for filtering and normalizing metabolomics data

**DOI:** 10.1101/387365

**Authors:** Courtney Schiffman, Lauren Petrick, Kelsi Perttula, Yukiko Yano, Henrik Carlsson, Todd Whitehead, Catherine Metayer, Josie Hayes, William M.B. Edmands, Stephen Rappaport, Sandrine Dudoit

## Abstract

**Introduction:** Untargeted metabolomics datasets contain large proportions of uninformative features and are affected by a variety of nuisance technical effects that can bias subsequent statistical analyses. Thus, there is a need for versatile and data-adaptive methods for filtering and normalizing data prior to investigating the underlying biological phenomena.

**Objectives:** Here, we propose and evaluate a data-adaptive pipeline for metabolomics data that are generated by liquid chromatography-mass spectrometry platforms.

**Methods:** Our data-adaptive pipeline includes novel methods for filtering features based on blank samples, proportions of missing values, and estimated intra-class correlation coefficients. It also incorporates a variant of k-nearest-neighbor imputation of missing values. Finally, we adapted an RNA-Seq approach and R package, *scone*, to select an appropriate normalization scheme for removing unwanted variation from metabolomics datasets.

**Results:** Using two metabolomics datasets that were generated in our laboratory from samples of human blood serum and neonatal blood spots, we compared our data-adaptive pipeline with a traditional filtering and normalization scheme. The data-adaptive approach outperformed the traditional pipeline in almost all metrics related to removal of unwanted variation and maintenance of biologically relevant signatures. The R code for running the data-adaptive pipeline is provided with an example dataset at https://github.com/courtneyschiffman/Data-adaptive-metabolomics.

**Conclusion:** Our proposed data-adaptive pipeline is intuitive and effectively reduces technical noise from untargeted metabolomics datasets. It is particularly relevant for interrogation of biological phenomena in data derived from complex matrices associated with biospecimens.

## 1 Introduction

Metabolomics represents the small-molecule phenotype that can be objectively and quantitatively measured in biofluids such as blood serum/plasma, urine, saliva, or tissue/cellular extracts (Chen et al., 2011; Escriva et al., 2017; Reinke et al., 2017; Want et al., 2013). Untargeted metabolomics studies allow researchers to characterize the totality of small molecules in a set of biospecimens and thereby discover metabolites that discriminate across phenotypes (Chen et al., 2011; Reinke et al., 2017; Scoville et al., 2018). Among the techniques employed for untargeted metabolomics, liquid chromatography-high resolution mass spectrometry (LC-HRMS) has become the analytical tool of choice due to its high sensitivity, simple sample preparation, and broad coverage of small molecules (Spicer et al., 2017; Want et al., 2013). However, many of the thousands of features detected by untargeted metabolomics are not biologically interesting because they represent multiple signals arising from the same analyte (adducts, isotopes, in-source fragmentation) and background signals from sample processing (Mahieu and Patti, 2017). Furthermore, metabolomic features are measured with multiple sources of technical variation, defined here as any random or fixed unwanted variability related to the measurement process. Examples of technical variation include perturbations in LCHRMS runs (e.g., variation in retention times and mass accuracy) and identifiable factors such as batch and run-order (De Livera et al., 2015; Herman et al., 2017; Patterson et al., 2016). Technical variation requires a set of preprocessing methods for filtering noise and normalizing features to adjust for biases prior to investigating the biological phenomena of interest. Furthermore, while many normalization techniques, such as linear regression (Ganna et al., 2016) and removal of unwanted variation (RUV) (Risso et al., 2014), have been proposed for use with metabolomics data, they have not been applied in a data-adaptive manner. In what follows, we present a series of steps (Fig. 1) representing a data-adaptive pipeline for filtering and normalizing untargeted metabolomics data prior to discovering common metabolites of potential interest, i.e., those that are present in most samples from contrasting populations. A data-adaptive pipeline is one which tailors filtering, imputation and normalization to the specific characteristics of a given data set, rather than using predefined methods. Using two untargeted LC-HRMS metabolomics datasets, we compare our approach to a traditional preprocessing pipeline and show that it more successfully uncovers meaningful biological phenomena.

**Fig. 1.**
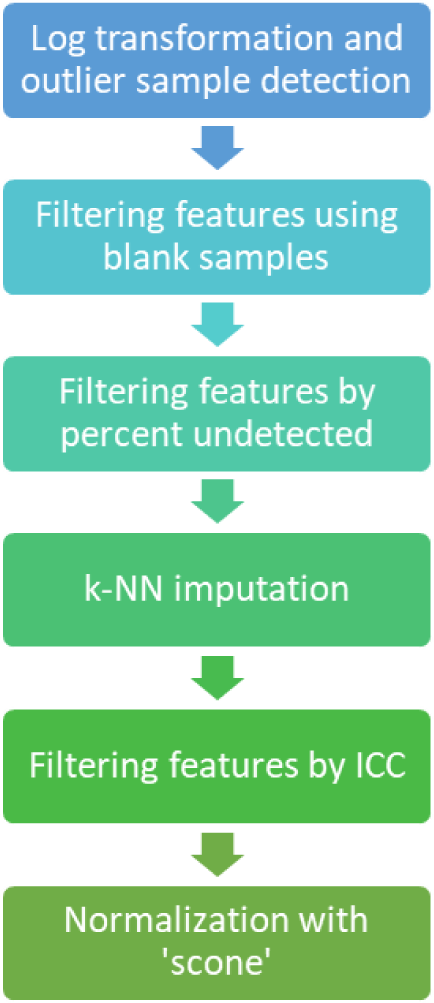
Flowchart of a data-adaptive filtering and normalization pipeline for untargeted metabolomics data.

## 2 Data-adaptive filtering and imputation

### 2.1 Log transformation and outlier detection

Feature abundances are transformed with the natural logarithm prior to filtering and normalization. Next, outlier samples are removed using the *pcaOutId* function from the *MetMSLine* R package (Edmands et al., 2015). This function uses a principle component score plot and Hotelling’s T-squared distribution ellipses to identify outliers. If the dataset contains multiple batches of samples, this procedure should be performed separately for each batch.

### 2.2 Filtering features based on blank samples

Blank control samples, which are obtained from the solvents and media used to prepare biological samples, can pinpoint background features that contribute to technical variation (Chen et al., 2011; Herman et al., 2017; Patterson et al., 2016; Want et al., 2013). A typical filtering method is to use a fold-change (biological signal/blank signal) cutoff to remove features that are not sufficiently abundant in biological samples (Chen et al., 2011). Rarely does the user examine the data to determine a suitable cutoff and a predetermined value of two to five is typically applied. We employ a data-adaptive procedure that takes into account the number of blank samples in which each feature is detected and then assigns cutoffs according to the background noise. If the dataset contains several batches, filtering is performed batch-wise.

We use a mean-difference plot (MD-plot) to visualize the relationship between feature abundances in the blank and biological samples and assess background noise (Fig. 2). The mean abundances of each feature in the biological and blank samples are calculated and the average of and difference between these two means are then plotted on the x- and y-axes, respectively. The horizontal zero-difference line (black line in Fig. 2) represents the cutoff between features having higher mean abundances in the blank samples and those having higher mean abundances in the biological samples. If there are *n* blank samples in a batch, then *n* + 1 clusters of features will typically be visually identifiable in the MD-plot, where cluster *i* = 0*,…, n* is composed of features that are detected in *i* blank samples (Fig. 2a). Filtering is performed separately for each cluster and features that are detected in none of the blank samples (cluster 0) are retained. For the remaining clusters, those that have non-uniform distributions of mean feature abundances are partitioned based on quantiles (20th, 40th, 60th, and 80th percentiles) of the empirical distribution of mean abundances (x-axis) (Fig. 2b). This ensures that each partition has the same number of features and that the features are uniformly distributed throughout the dynamic range. Within each partition, the empirical distribution of abundances below the zero-difference line is used to estimate the technical variation above that line and the absolute value (green lines in 2b) of the empirical first quartile of the negative mean differences (red lines in 2b) is used as a cutoff to remove uninformative features. Although it is possible for background signal to modify biological signal (e.g., via ion suppression), we do not consider this source of variability.

**Fig. 2.**
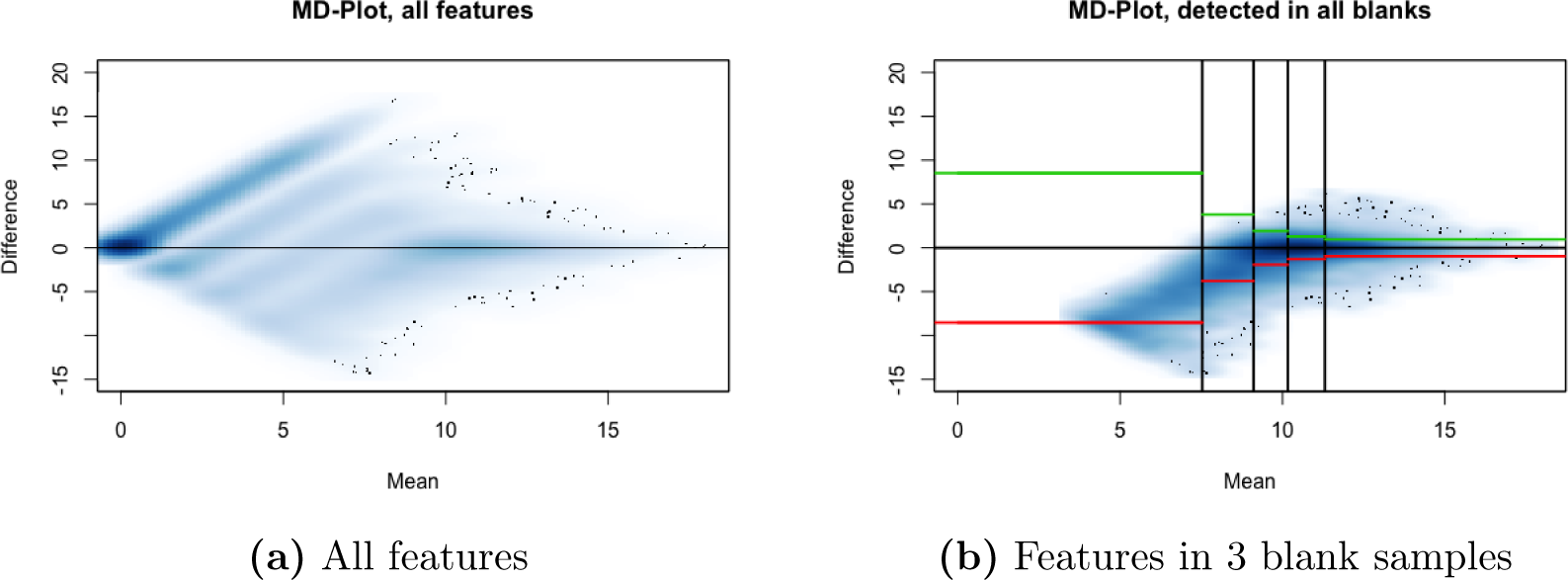
Filtering features based on blank sample abundances, NBS dataset. Mean-difference plots (MD-plots) of log abundances for the first batch of the neonatal blood spot (NBS) dataset described below (Petrick et al., 2017). **(a)** Four clusters of features are present: those detected in 3, 2, 1, and none of the blank samples. **(b)** Filtering thresholds for the cluster corresponding to features detected in all 3 blank samples.

### 2.3 Filtering features by percent missing

Low-abundance metabolomic features tend to have large numbers of undetected values across samples. A common way to handle undetected values is to use functions in standard metabolomics processing software, such as *fillPeaks* in the *XCMS* package, that impute values by integrating the background signal in the chromatographic region of the peak group (Ganna et al., 2016; Smith et al., 2006). However, in practice, some undetected values can remain unfilled even after using *fillPeaks*, thereby motivating removal of features with more than a predefined threshold of undetected values (e.g., 20%) (Reinke et al., 2017).

Because here we are focusing on common metabolites, we bypass the *fillPeaks* function and instead remove features with a high proportion of missing values. A data-adaptive cutoff is defined based on a Gaussian kernel density plot of the proportion of undetected values across all biological samples. Interestingly, both of the datasets evaluated in the results section, as well as other datasets collected in our laboratory, have density plots showing a similar bimodal pattern and suggesting a natural cutoff at 20% of undetected values for common features, retaining features near the first mode (Fig. 3a). It is not clear if this is a general or instrument-specific cutoff because all of our datasets were collected on the same analytical platform. Regardless, this density plot can easily be generated to determine an appropriate percent-missing cutoff for any dataset.

**Fig. 3.**
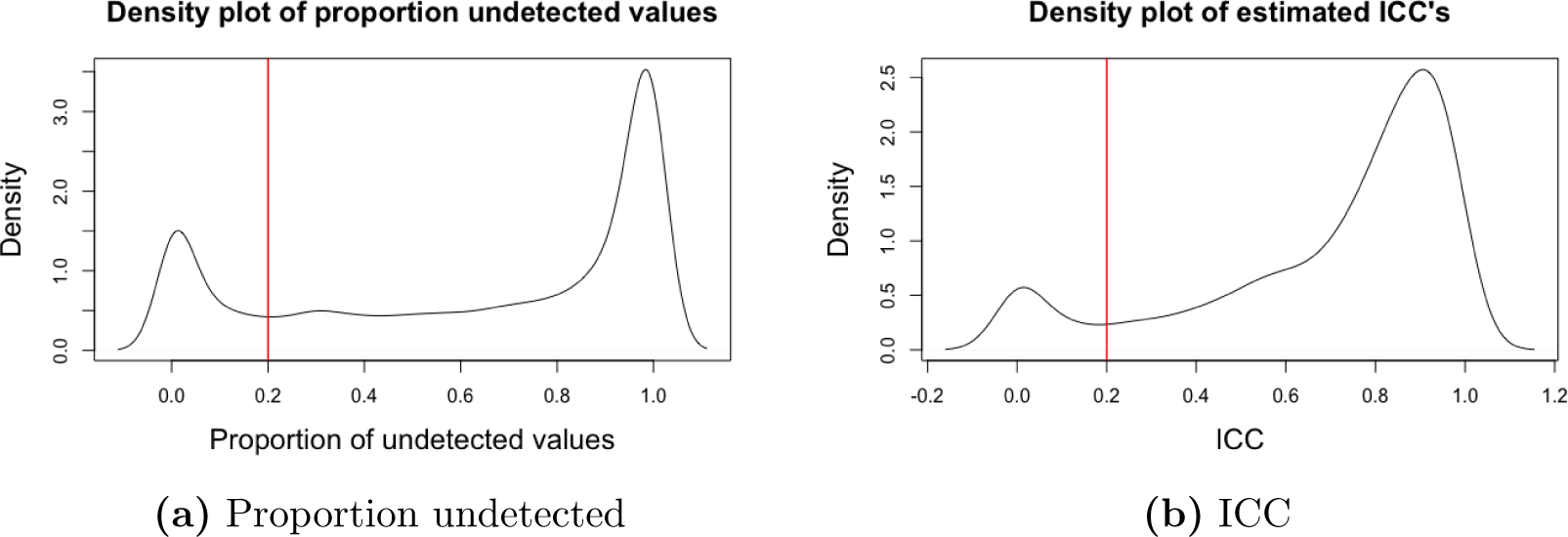
Filtering features by percent missing and ICC using Gaussian kernel density plots, NBS dataset. Gaussian kernel density plots of **(a)** the proportion of undetected values and **(b)** the estimated intra-class correlation coefficient (ICC) for each feature in the first batch of the NBS dataset (Petrick et al., 2017). Features with more than 20% undetected values or ICC less than 0.2 are removed.

### 2.4 Data imputation

The next step is to impute values for undetected features. Common imputation methods in metabolomics employ the limit of detection, the minimum abundance detected for each feature, k-nearest-neighbor (k-NN) imputation, or probabilistic principal component analysis (PPCA) (Xia and Wishart, 2016).

We prefer k-NN imputation because it is data-adaptive and has performed well for highdimensional and left-censored datasets (Do et al., 2018; Troyanskaya et al., 2001). To arrive at the imputed value, our pipeline seeks all *k* nearest-feature neighbors that are non-missing and averages their values (see Online Resource). Since features with undetected values tend to have low abundances, we select *k* as follows: randomly sample features (we used 100) among those with the lowest average abundances, set their abundances as missing in certain biological samples, impute these abundances using several values of *k*, and choose the *k* with the smallest mean squared error. We have found *k* = 5 to be a reasonable choice for untargeted metabolomics datasets in our laboratory.

### 2.5 Filtering features by ICC

Typically, the coefficient of variation (CV) across pooled quality control (QC) samples is calculated for each feature and those with a CV above a predetermined cutoff (e.g., 20-30%) are removed (Patterson et al., 2016; Reinke et al., 2017; Want et al., 2013). However, filtering features based solely on CV can remove some that are informative regarding the biology of interest. Instead, we propose examining the proportion of between-subject variation to total variation, otherwise known as the intraclass correlation coefficient (ICC) (Searle et al., 2006). A large ICC for a given feature would suggest that much of the total variation is due to biological variability and vice versa.

The method for estimation of the ICC employs the following one-way random effects model:

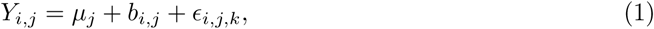

where *Y_i,j_* is the abundance of feature *j* in subject *i*, *µ_j_* is the overall average abundance of feature *j*, *b_i,j_* is a random effect for feature *j* in subject *i*, and *ϵ_i,j,k_* is a random error for replicate measurement *k* for feature *j* in subject *i*. The ICC is calculated by taking the ratio of the estimated variance of *b_i,j_* (between-subject variance) to the estimated variance of *b_i,j_* + *ϵ_i,j,k_* (total variance). If replicate specimens or LC-MS injections are analyzed for each subject, then application of Equation (1) is straightforward. However, since metabolomics data are often collected with single measurements of each biospecimen and employ repeated measurements of pooled QC samples to estimate precision, then Equation (1) can be fit by treating the pooled QC samples as repeated measures from a ‘pseudo-subject’.

As with the percent undetected values, a Gaussian kernel density plot of the estimated ICC values is used to select a cutoff for filtering features. For our two benchmarking datasets described in Section 4 and other datasets from our laboratory, we have observed a natural ICC cutoff of 0.2 (Fig. 3b), but an ICC density plot can be used to determine a suitable cutoff for a given dataset. If multiple batches are involved, the final feature list represents the intersection of features from all batches.

## 3 Data-adaptive normalization with scone

Each untargeted metabolomics dataset has a unique and often complex set of normalization requirements, such as adjustment for sample batch, run-order, processing and storage, dilution factors (e.g., creatinine for urine or potassium for dried blood spots), and known experimental contaminants (Pupillo et al., 2016). Therefore, to objectively select a suitable scheme for normalizing a given metabolomics dataset, we use the Bioconductor R package *scone* (Cole and Risso, 2017). While originally developed for single-cell RNA-Seq, *scone* implements the following normalization procedures that are immediately applicable to metabolomics data:

- global-scaling normalization, e.g., upper-quartile, *DESeq* (Anders and Huber, 2010), *TMM* (Robinson and Oshlack, 2010);
- full-quantile normalization;
- regression of scaled and logged feature abundances on
  – biological covariates of interest (e.g., disease status),
  – known factors of unwanted variation (e.g., batch),
  – estimated unknown factors of unwanted variation, as in RUV (Risso et al., 2014).

The *scone* package then evaluates each candidate scheme with metrics that gauge the removal of unwanted variation and retention of wanted variation. See Section 4 for detailed descriptions of the *scone* metrics.

## 4 Results and discussion

We compared our data-adaptive filtering and normalization pipeline to a more traditional pipeline, based on web servers like *MetaboAnalyst* (Xia and Wishart, 2016), popular software such as *XCMS* (Smith et al., 2006), and recent untargeted metabolomics studies (Cordeiro et al., 2018; De Livera et al., 2015; Ganna et al., 2016; Want et al., 2013). This traditional pipeline employs the following sequence:

1. The *fillPeaks* function from the *XCMS* software imputes most of the undetected values.
2. The *pcaOutid* function from the *metsmline* package removes outliers.
3. Features that remain undetected in more than 20% of the samples are removed.
4. Features with fold-changes (in biological vs. blank samples) less than five are removed.
5. Pooled QC samples are normalized by median scaling and features with CV greater than 30 % are removed.
6. The remaining missing values are imputed with half of the minimum detected value.
7. Biological samples are scaled by their median value for normalization purposes.
8. The abundance of each normalized feature is natural log-transformed.

We compared the data-adaptive and traditional pipelines with two untargeted metabolomics datasets generated in our laboratory with an LC-HRMS platform consisting of either an Agilent 1100 series or 1290 series LC coupled to an Agilent 6550 QToF mass spectrometer. One of these datasets represents the metabolomes of 4.7-mm punches from archived neonatal blood spots (NBS) of 400 control subjects that were obtained for the California Childhood Leukemia Study (Metayer et al., 2013) from the California birth registry, as described in Petrick et al. (2017). The second dataset contains the metabolomes of 122 serum samples from incident colorectal cancer (CRC) casecontrol pairs as described in Perttula et al. (2016). For comparison purposes, we define a binary biological factor of interest for each dataset, i.e., breastfeeding for the NBS (newborn infants are fed at least once prior to collection of NBS) and CRC diagnosis for the serum samples. To compare the two pipelines, we followed De Livera et al. (2015) and considered the presence and strength of association of positive control metabolites relative to the other features for either breast feeding (NBS) or case-control status (serum). Nominal *p*-value rankings were used to assess the relative ranks of the positive controls. However, since the two pipelines resulted in different sets of final features, the relative rankings should not be compared directly between the two pipelines. See the Online Resource for details of *p*-value calculations.

The filtered and normalized datasets from the two pipelines were compared based on several metrics borrowed from the *scone* package (referred to here as *biosil*, *batchsil*, *pamsil*, *expqccor*, and *expuvcor*). The metrics *biosil* and *batchsil* employ the average silhouette width of sample clusters (a measure of cluster ‘tightness’) to assess the degree to which the pipeline maintains the biological signal of interest (high *biosil* score) while reducing unwanted batch effects (low *batchsil* score). The metric *pamsil* assesses the preservation of sample clustering structure in terms of the average silhouette width of clusters obtained by applying partitioning around medoids (PAM) (Kaufman and Rousseeuw, 1990) to the first three principal components of normalized abundances (a large *pamsil* is preferred). The metrics *expqccor* and *expuvcor* represent *R*^2^ values from regressions of the first three principal components of the final feature abundances on the first *k* principal components of the known factors of unwanted variation or the first *k* estimated factors of unwanted variation, respectively (lower scores for *expqccor* and *expuvcor* are preferred). See the Online Resource for more details on the *scone* metrics.

Appropriately filtered and normalized datasets from a given pipeline should have relative log abundances (RLA) centered at zero with small variability (De Livera et al., 2015). Therefore, four additional metrics (*rlemed1*, *rlemed0*, *rleiqr1*, and *rleiqr0*) were used to measure the median RLA and variance of the inter-quartile range within biological groups of interest. When data are suitably normalized, the distribution of *p*-values from testing associations of feature abundances with a given biological factor should be mostly uniform, with a peak near zero for associated features (De Livera et al., 2015). Thus, we assessed the uniformity of the distribution of *p*-values using quantile-quantile plots (QQ-plots). Visualization of the first and second principal components of the final filtered and normalized abundances were also used to assess the removal of unwanted variation.

### 4.1 Metabolomics of neonatal blood spots

Over 60,000 features were initially measured in each of 400 NBS samples that were analyzed in four batches (along with blank and QC samples). Because the biological factor of interest is breastfeeding, two fatty acids that are abundant in breast milk, i.e., palmitoleic acid and docosahexaenoic acid (DHA) (Chuang et al., 2013; Gardner et al., 2017), were used as positive controls, selected prior to any analysis. Known sources of technical variation in these samples include NBS age, potassium level (a measure of hematocrit), run-order, and batch (De Kesel et al., 2014; Pupillo et al., 2016).

#### 4.1.1 Traditional pipeline

For the traditional pipeline, the vast majority of features passed the maximum 20% missing values cutoff (Fig. 5a) because the *fillPeaks* function imputed most undetected values. However, because features with large proportions of imputed values had low average abundances, filtering by a foldchange cutoff of five reduced the number of features (Fig. 5a) from over 60,000 to 8,726, and further filtering by a CV cutoff of 30% resulted in a final set of 1,349 features. The positive control metabolite palmitoleic acid was not included in this final dataset because its CV of 36% exceeded the preassigned cutoff of 30%. The other positive control (DHA) was present in the final dataset at an average level that was 1.3 times higher in breastfed babies and ranked 32nd among all features in terms of strength of association with breastfeeding.

#### 4.1.2 Data-adaptive pipeline

For the data-adaptive pipeline, filtering by blank samples reduced the number of features from 60,000 to 25,000, subsequent filtering by a cutoff of 20% missing in each batch (Fig. 3a) resulted in 1,607 features, and the final filtering by an ICC cutoff of 0.2 resulted in a total of 1,070 features (Fig. 3b). Palmitoleic acid had the eighth smallest *p*-value among all features when testing for association with breastfeeding and was present at an average level that was 1.4 times higher in breastfed infants. DHA was also present in the final dataset and ranked 97th among all features for association with breastfeeding, with an average level that was 1.2 times higher in breastfed infants. The top normalization scheme ranked by *scone* included no global-scaling and regression-based adjustment for six estimated factors of unwanted variation.

#### 4.1.3 Pipeline comparison

The data-adaptive pipeline outperformed the traditional pipeline for eight out of the nine *scone* and RLA metrics, which assessed the degree to which unwanted variation from batch, run-order, spot age, and potassium level was removed, and thereby enhanced associations with breastfeeding status (see Online Resource). Only the *scone* metric *pamsil*, a measure of sample heterogeneity based on PAM clustering, was ranked more highly for the traditional pipeline. Although the QQ-plots were similar for both pipelines, the *p*-value distribution was more uniform for the data-adaptive pipeline, with a departure near zero related to features associated with breastfeeding (see Online Resource). Principal component plots also show that the data-adaptive pipeline was more effective at removing unwanted variation by filtering and normalization (Fig. 4a,b). Both positive-control metabolites were retained by the data-adaptive pipeline and were ranked relatively highly (i.e., among the smallest *p*-values), whereas the traditional pipeline only retained DHA that was also ranked relatively highly.

**Fig. 4.**
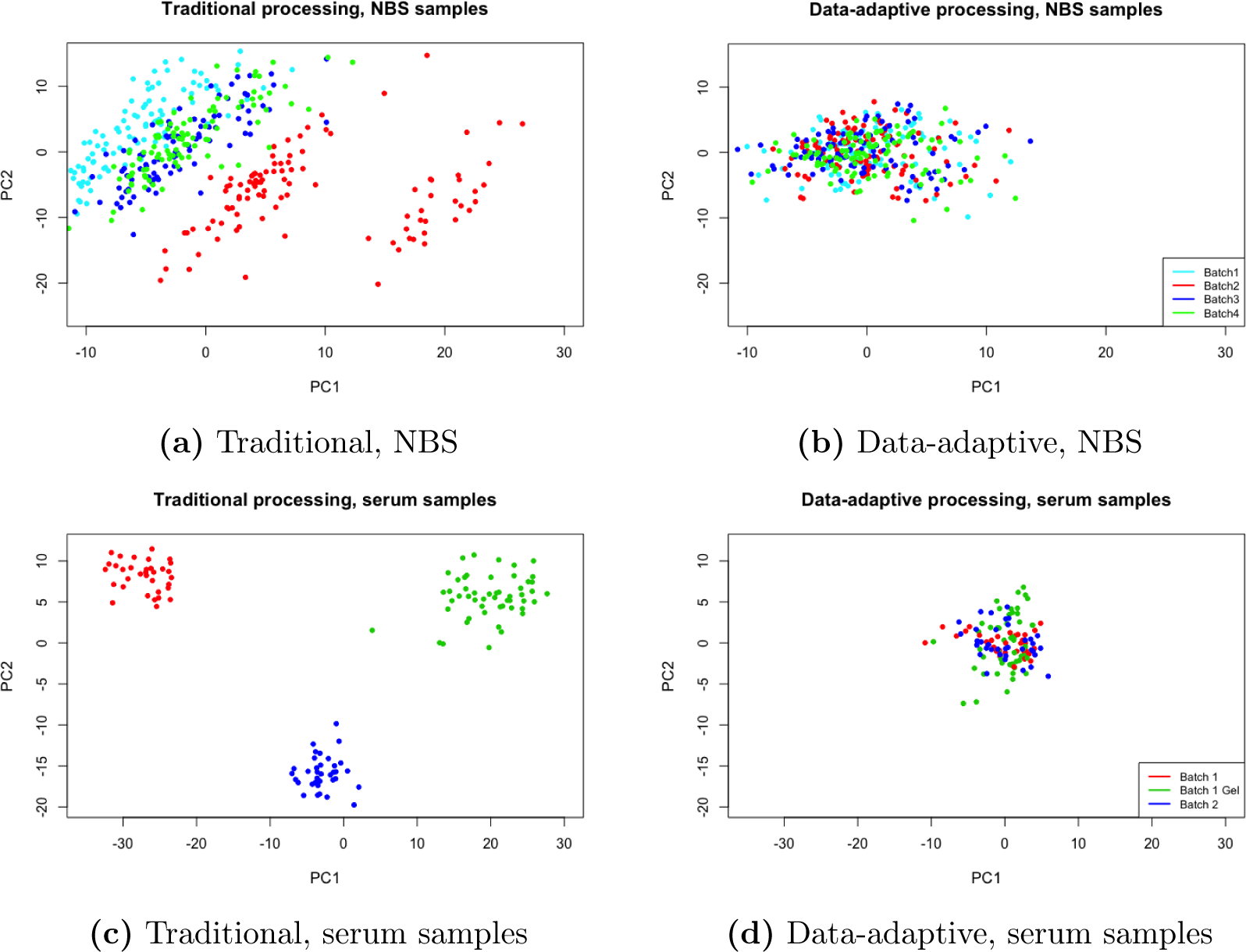
Traditional and data-adaptive filtering and normalization, NBS and serum datasets. Principal component plots for the NBS (Petrick et al., 2017) and serum (Perttula et al., 2016) datasets after traditional and data-adaptive filtering and normalization. Samples are color-coded by batch.

### 4.2 Metabolomics of serum samples

Over 20,000 features were detected in the 122 serum samples (61 incident cases, 61 controls) that were analyzed in two batches (Perttula et al., 2016). Fifty-two of the samples in the first batch were contaminated by additives that resulted from gelled serum, thereby introducing an additional known source of technical variation into the data. Sample age was not considered as a source of variation, because all specimens were collected during a small time window and were stored in liquid nitrogen soon after collection. As positive controls for CRC, we used eight ultra-long-chain fatty acids (ULCFA 446, 448, 466, 468, 492, 494, 538, 594), that had previously been shown to be present at lower levels in CRC cases than controls in several cross-sectional studies as well as in this set of samples (summarized by Perttula et al. (2016)).

#### 4.2.1 Traditional pipeline

In the traditional filtering pipeline, most features (20,830) again passed the 20% missing cutoff (Fig. 5a). However, filtering by a fold-change of five reduced the number of features to 13,391 and filtering by CV less than 30% further reduced the number to 2,536. Five of the eight positive controls (ULCFA 446, 466, 468, 492, 494) were removed because their CV values were above 30% (34.6%, 41.6%, 34.5%, 42.5%, 33.2%, respectively). The three remaining positive controls (ULCFA 448, 538, 594) ranked 1210th, 1039th, and 2045th in terms of *p*-values, respectively, with case-control fold-changes of 0.87, 0.85, and 0.94.

**Fig. 5.**
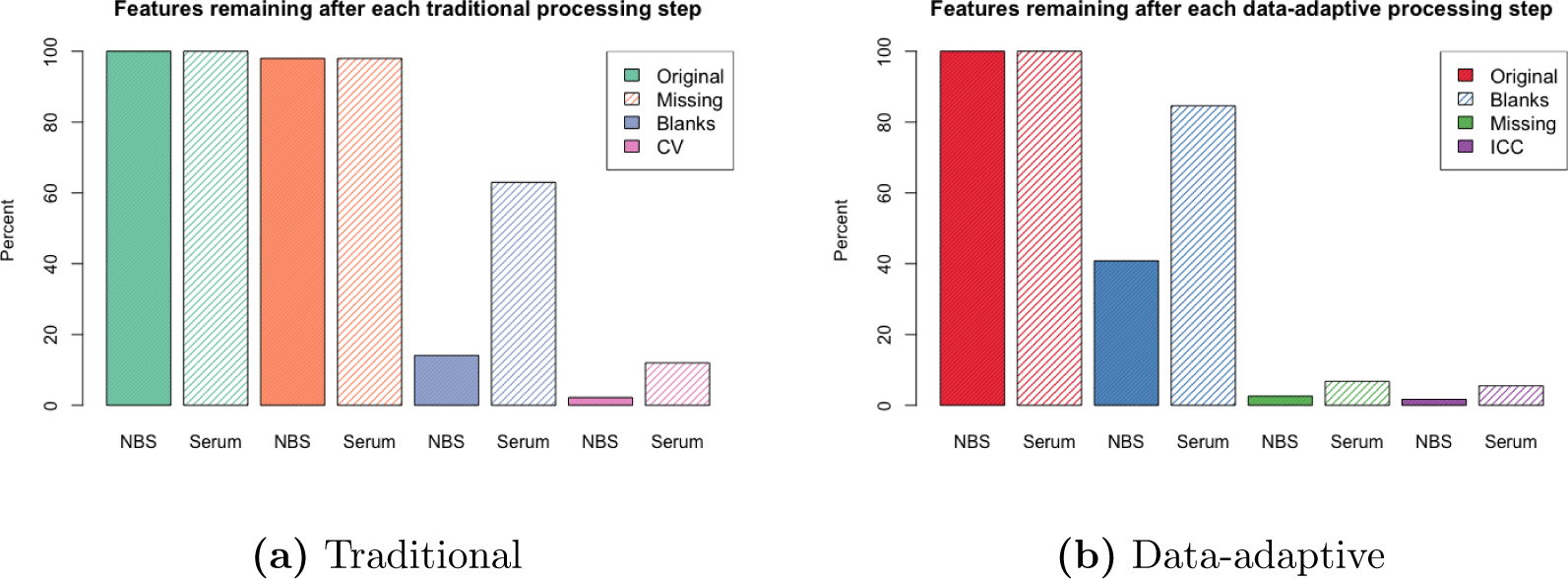
Traditional and data-adaptive filtering steps, NBS and serum datasets. The percentage of total features retained in each step of the **(a)** traditional and **(b)** data-adaptive filtering pipelines, for the NBS (Petrick et al., 2017) and serum (Perttula et al., 2016) datasets.

#### 4.2.2 Data-adaptive pipeline

In the data-adaptive pipeline, filtering based on blank abundances left 17,692 features; the majority of these (around 17,000) were then removed because more than 20% of their values were missing (Fig. 5b). Filtering by ICC values removed an additional 77 features, leaving 378 in the final dataset. The top normalization scheme ranked by *scone* included *DESeq* global scaling and regression-based adjustment for batch, gel contamination, and run-order. Four positive controls (ULCFA 446, 468, 538, 594) were retained in the final dataset, with *p*-values ranking 22nd, 2nd, 12th, and 187th and case-control fold-changes of 0.75, 0.73, 0.79, and 0.90, respectively. Three of the positive controls (ULCFA 466, 492, 494) were removed because they contained over 80% missing values in the second batch and one was removed with an ICC of 0.18 (ULCFA 448).

#### 4.2.3 Pipeline comparison

The data-adaptive pipeline outperformed the traditional pipeline for six of the nine metrics concerning removal of unwanted variation (e.g., batch, gel contamination, run-order) and characteristics of RLA distributions, and tied with the traditional pipeline for the *rleiqr1* metric (see Online Resource). The traditional processing had higher scores for only the *biosil* and *pamsil* metrics. Although QQ-plots of *p*-values for both pipelines showed heterogeneous distributions, the lack of homogeneity was more severe for the traditional pipeline (see Online Resource). Principal component plots showed pronounced batch and gel-contamination effects in the dataset from the traditional pipeline, but not in that from the data-adaptive pipeline (Fig. 4c,d). All three positive controls retained by the traditional pipeline had relatively low ranking *p*-values (1040th, 1199th, and 2045th), whereas three of the positive controls retained by the data-adaptive pipeline had high ranking *p*-values (2nd, 12th, and 22nd). ULCFAs 468 and 446, which ranked 2nd and 22nd in the data-adaptive pipeline, were removed by the traditional pipeline because their CV were above 30%. ULCFA 448 was retained by the traditional pipeline but not the data-adaptive pipeline because it had a small ICC value (0.18) and unsurprisingly ranked 1210th in terms of *p*-values.

### 4.3 Discussion

Comparisons between our data-adaptive pipeline and a more traditional approach for preprocessing metabolomics data suggest that the data-adaptive pipeline offers several advantages. Although both pipelines have ‘high-impact’ filtering steps that remove large proportions of data (Fig. 5), the traditional pipeline is more likely to remove biologically informative features. For example, of the roughly 21,000 features in the NBS dataset that were not detected in any of the blank samples, only 6,000 were retained by the traditional pipeline after fold-change filtering with a cutoff of five (Fig. 5a). This massive loss of potentially interesting features would only have been partially rectified by reducing the fold-change cutoff from five to two or three because the median fold-change for these 21,000 features was 2.43. In contrast, the data-adaptive pipeline necessarily retained all features that were not detected in any of the blank samples (Fig. 5b).

Filtering by the coefficient of variation (a measure of relative variability) in the traditional pipeline also greatly reduced the number of potentially important features (Fig. 5a). Indeed, a CV cutoff of 30% removed positive controls from both datasets (palmitoleic acid and ULCFA 446, 466, 468, 492, 494 for the NBS and serum samples, respectively). In contrast, data-adaptive filtering by the proportion of between-subject variation to total variation (ICC) removed fewer features (Fig. 5 b) while retaining both positive controls in the NBS samples and four of the eight ULCFAs in the serum samples. We interpret this to mean that some imprecisely-measured features differed substantially across subjects and, therefore, held meaningful biological information.

Besides the high-impact filtering steps, another notable difference between the two pipelines is the inclusion of the *scone* framework to objectively normalize features abundances prior to statistical analyses such as feature selection or sample-class prediction. As is typical with untargeted metabolomics, both datasets contained considerable technical variation due to long sample runs, batch effects, contamination, etc. Such complex sources of unwanted variation require normalization schemes that are more aggressive than global scaling as applied by the traditional pipeline (Fig. 4a,c), and demand a framework for objectively comparing different approaches. Indeed, for both datasets, *scone* motivated informed normalization that outperformed the traditional pipeline for all metrics involving removal of unwanted variation (see Online Resource).

The handling of undetected features also has important implications. Whereas the traditional pipeline employed the *fillPeaks* function to impute baseline abundances for undetected samples, the data-adaptive pipeline employed k-NN to impute abundances for features with fewer than 20% missing values. By invoking *fillPeaks* at the outset, the traditional pipeline inflates the percentage of imputed values in any final list of features beyond an arbitrary cutoff of say 20%, a cutoff we found to be reasonable for our two datasets (Fig. 3). For example, after applying the traditional pipeline to the NBS dataset, half (675) of the final 1,349 features had more than 34% values imputed by *fillPeaks* in at least one of the four batches compared to only nine of the 1,070 final features from the data-adaptive pipeline having 20% values imputed by k-NN. Furthermore, we have observed that metabolomic features with large proportions of undetected values tend to have poorly integrated peaks and low mean abundances that can complicate subsequent analyses. By removing such features, the data-adaptive pipeline arguably provides more robust data for statistical analyses and enhances the prospects for annotation (see Online Resource).

The data-adaptive pipeline can be modified to suit the specific goals of each untargeted metabolomics study, such as the discovery of rare metabolites. In such a study, the minimum number of samples required for a peak group to be present in (e.g., the *minfrac* parameter in the *XCMS* software) can be lowered, the absolute value of the empirical median (rather than the first quartile) can be used as a cutoff when filtering by blank sample abundances, and a higher proportion of undetected values can be allowed (e.g., 80% in Fig. 3a). A step can also be added to the data-adaptive pipeline to retain features with missing values that are differentially expressed across biological groups of interest, e.g., features that are mostly missing among controls but not incident cases. These modifications to the data-adaptive pipeline would result in considerably more features, but with higher proportions of missing values that can complicate subsequent statistical analyses.

## 5 Conclusions

Untargeted metabolomics generates data with varied sources of technical noise – some unique to a given study – that benefit from data-adaptive preprocessing prior to statistical analyses. Here, we provide one such pipeline and demonstrate that it removes unwanted variation while maintaining biological variability in metabolomic features. When compared to a traditional metabolomics pipeline, the data-adaptive approach performed better for essentially all objective criteria including *scone* metrics, RLA metrics, lack of clustering of samples by unwanted factors in PC plots, and retention of positive-control metabolites.

## Acknowledgements

We gratefully acknowledge the assistance of Agilent Technologies (Santa Clara, CA, USA) for the loan of the high-resolution mass spectrometer that was used to generate these two datasets.

## Author contributions

CS developed the pipeline,performed all filtering, normalization, and statistical analyses, and wrote the manuscript. LP helped to develop the pipeline, collected the NBS dataset, and assisted with writing the manuscript. SD and SR supervised data collection and pipeline development and contributed to writing the manuscript. KP collected the serum sample dataset. HC annotated the palmitoleic acid positive control. YY provided potassium measurements for normalization. TW and CM contributed the NBS samples. WMBE helped collect both benchmark datasets. JH processed the raw data for the serum dataset.

## Compliance with ethical requirements

### Research involving animal and human rights

The datasets used for benchmarking were obtained from two previously published studies. The study was approved by the University of California Committee for the Protection of Human Subjects, the California Health and Human Services Agency Committee for the Protection of Human Subjects, and the institutional review boards of all participating hospitals. The biospecimens (neonatal blood specimens) and corresponding data used in this study were obtained from the California Biobank Program, (SIS request number(s) 26), Section 6555(b), 17 CCR. The California Department of Public Health is not responsible for the results or conclusions drawn by the authors of this publication.

### Informed Consent

Both investigations obtained biospecimens from human subjects with informed written consent under protocols that had been approved by institutional review boards from all participating institutions. Written informed consent to participate in childhood leukemia research was obtained from the parents of all study subjects from the California Childhood Leukemia Study.

### Funding

This work was supported by the National Institute for Environmental Health Sciences (NIEHS) of the U.S. National Institutes of Health and the U.S. Environmental Protection Agency (USEPA) through grants to the Center for Integrative Research on Childhood Leukemia and the Environment (NIEHS grants P01 ES018172 and P50ES018172 and USEPA grants RD83451101 and RD83615901), by the California Childhood Leukemia Study (NIEHS grants R01ES009137 and P42ES004705), and by grant agreement 308610-FP7 from the European Commission (Project Exposomics). NIEHS and USEPA are not responsible for the results or conclusions drawn by the authors of this publication.

### Software availability

The pipeline developed in this study and an example dataset for running it are available via https://github.com/courtneyschiffman/Data-adaptive-metabolomics.

